# Roles of hostplant availability and quality for the distribution and climate change response of a dietary specialist herbivore

**DOI:** 10.1101/2020.12.03.410225

**Authors:** A. Nalleli Carvajal Acosta, Kailen Mooney

**Affiliations:** Department of Ecology and Evolutionary Biology, University of California-Irvine, Irvine, California, United States of America

**Keywords:** *Danaus plexippus*, monarch, species distribution models, MaxEnt, climate change, bioclimatic models, hostplant quality, Milkweeds, *Asclepias*, biotic interactions, specialized herbivores

## Abstract

Species distributions are recognized to be driven by abiotic factors, but the importance of biotic interactions that provide critical resources is less well understood, especially with respect to variation in critical resource quality. Disentangling the relative importance of these factors – abiotic environment, presence of critical resources and their quality-may be critical to predicting species response to climate change. We used species distribution models (SDMs) to address these questions for the western monarch butterfly (*Danaus plexippus*), a species that obligately feeds upon plants in the genus *Asclepias*, and for which hostplant quality in this region varies among species by an order of magnitude. We modeled the distribution of 24 *Asclepias* species to develop and compare three monarch distribution models with increasing levels of ecological complexity: (i) a null model using only environmental factors (a climate envelope model), (ii) a model using environmental factors and *Asclepias* spp. distribution, (iii) and a model using environmental factors and *Asclepias* spp. distribution weighted by hostplant quality assessed through a greenhouse bioassays of larval performance. *Asclepias* models predicted that half of the *Asclepias* spp. will both expand their ranges and shift their distribution towards higher latitudes while half will contract within the study region. Our performance analysis of monarch models revealed that the climate envelope model was the poorest performing. Adding hostplant distribution produced the best performing model, while accounting for hostplant quality did not improved model performance. The climate envelope model estimated more restrictive contemporary and future monarch ranges compared to both hostplants models. Although all three models predicted future monarch range expansions, the projected future distributions varied among models. The climate envelope model predicted range expansions along the Pacific coast and contractions inland while hostplants models predicted range expansions in both of these regions and, as a result, estimated 14 and19% increases in distribution relative to the climate envelope model, respectively. These results suggest that information on biotic interactions that provide critical resources is needed to predict future species distributions, but that variation in the quality of those critical resources may be of secondary importance.

## Introduction

Climate change is expected to alter the distribution of most species (Parmesan et al. 1999, Crozier 2004, Bellard et al. 2012, Pauli et al. 2012) with many already experiencing range contractions or facing extinctions (Sekercioglu et al. 2008, La Sorte and Jetz 2010, Bellard et al. 2012, Pauli et al. 2012). Understanding the underlying drivers is key to predicting such distributional response and also critical if we are to mitigate these impacts. Species distributions are presumed to be driven most strongly by abiotic factors, but biotic interactions can also play a key role (Guisan and Thuiller 2005). Because species often respond differently to abiotic stress (Schweiger et al. 2008, Van der Putten et al. 2010), producing accurate predictions necessitates that we also account for climate change effects on interacting species. This is especially true for species that engage in obligate interactions, as they depend on a few or even a single species to survive, and such species may not be available in all areas that are otherwise climatically suitable (Schweiger et al. 2008).

Herbivorous insects – the majority of multi-cellular species on earth (Lewinsohn et al. 2005) – are highly host-specific; thus, their response to climate change will likely depend fundamentally on the responses of the plants upon which they are obligately dependent. Indeed, most herbivorous insects feed on a single or a few plant families (Bernays 1989, Forister et al. 2015) with fewer than 10% feeding on plants belonging to more than three families (Price 1983). Furthermore, it has long been recognized that hostplants demonstrate considerable intra- and inter-specific variation in their resource quality to herbivores, and that resource quality is often heterogeneously distributed across landscapes (Denno and McClurc 1983, Hunter et al. 1992). Intra- and inter-specific variation in host-quality can have large effects on herbivore performance (Singer et al. 2012) and may also play a significant role in determining the spatial distribution of host-specific herbivorous insects at local scales (Memmott et al. 1995, Mcmillin and Wagner 1998, Egan and Ott 2007). However, the role of hostplant quality as a driver of species distribution at large spatial scales, and its implications for herbivore’s distributional response to climate change, are largely unknown.

In this study we investigated the importance of hostplant distribution and quality as drivers of herbivore contemporary distribution and response to projected future climate change. We use Species Distribution Models (SDMs), statistical tools that combine observations of species occurrences with environmental covariates to estimate species distributions. These models identify the factors driving contemporary species ranges and can also infer species response to climate change based on projections for how those driving factors will change in the future (Elith and Leathwick 2009). SDMs have most often assumed that species distributions are defined by environmental factors alone. This so-called “climate envelope approach” are based on the Eltonian noise hypothesis, which posits that biotic interactions may be a major driver of abundance at smaller spatial resolutions, but at larger and coarser spatial resolutions the effects of biotic interactions may average out, leaving abiotic factors as the principal drivers (Guisan and Thuiller 2005, Soberon and Nakamura 2009, Elith and Leathwick 2009). Yet recent modelling studies have identified biotic factors as important drivers of species distributions (Dilts et al. n.d., Araújo and Luoto 2007, Preston et al. 2008, Schweiger et al. 2008, de Araújo et al. 2014, Fraterrigo et al. 2014, Lemoine 2015, da Cunha et al. 2018) and SDMs predictions for species response to climate change have yielded contrasting results based upon whether or not biotic factors are included (Preston et al. 2008, Schweiger et al. 2008, Lemoine 2015). Accordingly, climate envelope modeling may accurately define the potential niche of a species, but the realized niche – defined in part by species interactions – may be substantially smaller.

Our aim in this study was to assess the importance of hostplant distribution and quality for driving contemporary and future distributions of dietary specialist herbivores. To do so, we studied the monarch butterfly (*Danaus plexippus*, Lepidoptera: Nymphalidae), the larvae of which feed exclusively from plants in the *Asclepias* genus which varies greatly among species in herbivore-defenses traits, nutrient content, and overall host quality (Agrawal and Fishbein 2006, Pocius et al. 2017). Monarchs are well known for their migratory and overwintering behavior (Pelton et al. 2019), and three previous studies have modelled their distribution. Lemoine (Lemoine 2015) accounted for hostplant distribution in the eastern monarch population response to climate change, predicting a poleward range expansion facilitated by *Asclepias* range expansions. Steven and Frey (Stevens and Frey 2010), and more recently Dilts *et al* (Dilts et al. n.d.), examined the role of hostplant availability and climate in determining the contemporary western monarch distribution and their breeding grounds, again demonstrating the importance of hostplants availability.

In the present study, we investigate the role of both hostplant distribution and quality in driving contemporary and future distributions of the western monarch population. To do so, we compared the performance of three species distribution models. In order of increasing complexity, these three models were: (i) a model using only climatic variables as predictors (hereafter, climate envelope model); (ii) a model using climatic variables and *Asclepias* distribution as predictors (hereafter, hostplant-presence model); and (iii) a model that included climatic variables, hostplants distribution and hostplant quality, which varied 10-fold among species as assessed through bioassays of larval performance (hereafter, hostplant quality model). We compared model performances and identified the variables determining the distribution of the western monarch breeding ranges. These models were then used to project and estimate changes in their distribution. Our study adds to past studies of this species and represents the first to estimate the future breeding range of the western monarch population. More broadly, this study is, to our knowledge, the first to explicitly test for the importance of hostplant quality of an obligate resource in driving species contemporary and future distribution.

## Materials and methods

### Study System

Monarch butterflies occur world-wide and, in their larval stage, feed exclusively from plants in the milkweed family (*Asclepias*, Apocyneceae: Asclepiadaceae). In North America, there are two migratory populations that breed east and west of the Rocky Mountains, with each of these regions being populated by multiple and largely unique sets of hostplant species (Ladner and Altizer 2005). Despite its dramatic population decline (Pelton et al. 2019), the western monarch population has been considerably understudied in comparison to the largest eastern population and we know little about how this population will be affected by climate change.

Western monarchs breed west of the Rocky Mountains and overwinter along the Pacific coast from Bodega Bay in northern California and as far south as Ensenada, Baja California, Mexico (Stevens and Frey 2010). During the spring, monarchs leave their overwintering sites and disperse throughout the western U.S. where they breed continuously during the summer. In the fall, adult monarchs return to their overwintering grounds (Pelton et al. 2019). Within North America, monarchs have been recorded feeding on 27 different plant species in the genus *Asclepias* (Ladner and Altizer 2005); however, adult females may oviposit in any available *Asclepias* species. Thus, monarchs may utilize multiple *Asclepias* species throughout their migratory paths.

The genus *Asclepias*, commonly known as milkweeds, consists of over 140 different species of which 130 are endemic to North America (Agrawal and Konno 2009). Milkweeds vary in their herbivore defensive strategies, which variously include combinations of cardenolides, latex, and trichomes, among others traits (Agrawal and Fishbein 2006). Interspecific variation in the quantity of plant defenses (Agrawal and Fishbein 2006) and nutrient content (Pocius et al. 2017) have been associated with monarch larval mass, developmental rate, and early instar survival (Zalucki et al. 2001). In this sense, the quality of the *Asclepias* species may be important in determining monarch distributions.

### Data Collection

#### Occurrence data

We retrieved monarch and milkweed records for the United States using R Studio (R Studio Team 2015) from multiple open source databases using the R packages SPOCC, Ecoengine, rbison (Chamberlain et al. 2014, Karthik 2014, Chamberlain 2019) and by accessing species occurrences directly from GBIF and iNaturalist databases (“GBIF Occurrence Download” 2019, “Naturalist [online]. Website” 2019). For monarchs, we only selected eggs and larval records because they provide a direct index for the location of the monarchs breeding grounds as opposed to adult records which may only indicate the migratory path. Additional monarch larval records were provided by the Monarch Larvae Monitor Program (MLMP) (Ries and Oberhauser 2015).

The occurrence data archived in open source databases originates mainly from citizen scientist sightings and some from herbarium records. As opposed to formal survey methods, this type of data has some limitations such as sampling biases, potential misidentification and coordinate inaccuracies, and lack species absence records. We controlled for these limitations whenever possible. For example, when permitted, we used filters that only retrieved records confirmed by experts and/or records classified as of research quality and spatial filtering to control for sampling biases.

To focus on the western monarch population, we selected Milkweeds and monarch larval records from states corresponding to this region: California, Nevada, Colorado, Washington, New Mexico, Arizona, Utah, Oregon, and Idaho. After removing duplicate records, incorrect (i.e. over oceans) or inaccurate coordinates (>1000 meters uncertainty) and observations, the final databases included 7,941 Milkweed records for 51 species (Data S1), and 904 monarch larval records (Data S2). *A. fascicularis* and *A. speciosa* were the most common species with 22% (2,541) and 12% (1,404) of total Milkweed records, respectively.

#### Environmental data and climate projections

Contemporary environmental bioclimatic variables and projections for the year 2070 were downloaded in R from the WorldClim website (Fick and Hijmans 2017) at 30-sec (approximately 1-km^2^) grid cells, the finest spatial resolution available. The current bioclimatic variables represent averages of a 50-year period from 1950 to 2000. Climate change projections for the year 2070 represent averages of a 30-year period from 2061 to 2080 based on the Hadley Centre Global Environmental Model, version 2, Earth System (HadGEM2-ES) model. The HadGEM2-ES model is recommended for ecological modeling as it accounts for ecologically-meaningful processes such as dynamic vegetation cover (The HadGEM2 Development Team: G. M. Martin et al. 2011). These projections are based on Representative Concentration Pathway (RCP) 8.5. The RCP 8.5 represents the worst-case scenario for greenhouse gas (GHG) concentrations, assuming that GHG emissions will continue to increase after the 21^st^ century in contrast to other scenarios that assume GHG will remain stable or will decline after the 21^st^ century (Collins et al. n.d.). While a comparison of different projections for future climate would provide a more nuanced prediction for the future distributions of milkweeds and monarchs, using this single scenario met our primary purpose of evaluating the importance of host plant information in predicting specialist herbivore distributions.

Environmental layers were cropped to include the states corresponding to range of the western monarch population. To reduce multicollinearity among variables, we removed highly correlated variables based on their Pearson correlation coefficients using a pairwise correlations approach following Dormann *et al*. (Dormann et al. 2013) but with a less restrictive threshold of 0.85 as in Elith *et al*. (Elith et al. 2006). We first removed variables that were correlated with multiple variables and, when only two variables were correlated, we selected the variable that was less statistically derived. This process yielded 11 environmental predictors (Table 1).

**Table 1.** Selected environmental variables

| **Worldclim Code** | **Environmental Variable\*** |
| --- | --- |
| **Bio1** | Annual Mean Temperature |
| **Bio5** | Max Temperature of Warmest Month |
| **Bio6** | Minimum Temperature of Coldest Month |
| **Bio7** | Temperature Annual Range |
| **Bio8** | Mean Temperature of Wettest Quarter |
| **Bio9** | Mean Temperature of Driest Quarter |
| **Bio12** | Annual Precipitation |
| **Bio15** | Precipitation Seasonality |
| **Bio17** | Precipitation of Driest Quarter |
| **Bio18** | Precipitation of Warmest Quarter |
| **Bio19** | Precipitation of Coldest Quarter |

\*Selected environmental variables with Pearson correlation coefficient of 0.85 or lower.

### Species distribution modeling

Because species occurrences in these datasets are available in the form of presence-only records, we used the maximum entropy method (hereafter MaxEnt) (Phillips et al. 2006) to model the current and future distribution of *Asclepias* and monarch breeding ranges. The MaxEnt algorithm is a presence-background modeling tool based on Bayesian and maximum likelihood statistics (Elith et al. 2011). To estimate the probability of distribution of a species, MaxEnt uses species presence records and a set of environmental predictors (e.g. precipitation, temperature) across a pre-defined landscape that is divided into grid cells. From this landscape, background points are randomly selected to represent the species environmental domain or background environment. MaxEnt estimates the relative probability of occurrence for each grid cell by maximizing the similarity between the environmental conditions of presence records and that of the background environment, while constraining the prediction to have the same mean as the presence records. The relative probabilities (raw output) are transformed to probability of occurrence using post-logistic transformation (logistic output). Here we report the logistic output which assigns a probability of presence between 0 and 1 to each grid cell, assuming that typical presence localities have a probability of presence of 0.5. See Elith *et al*. (Elith et al. 2011) for a comprehensive statistical explanation of MaxEnt.

Data collection, data processing, and modeling were performed in R studio (R Studio Team 2015). Species distribution modeling was executed in MaxEnt using the ‘dismo’ package (Hijmans et al. 2011).

#### Asclepias models

We developed models for individual *Asclepias* species and estimated their distributions within an area restricted to the study region; therefore, our *Asclepias* ranges do not represent their full distributions but only represent hostplant availability for the western monarch. *Asclepias* species were modelled separately because their distributions may be delimited by distinct environmental factors. We discarded records identified at the genus level and species with fewer than 40 records as this limited number of observations would not allow for an accurate estimation of their distributions. To correct for potential sampling biases, we used a spatial filtering approach which consists on randomly selecting one record per grid cell of a specified size (Kramer-Schadt et al. 2013). Spatial filtering was performed individually for each *Asclepias* species. This allowed us to retain records for multiple species co-occurring within a single grid cell as well as selecting the optimal spatial resolution that maximizes sample size while correcting for sampling biases. For example, species with a limited distribution (e.g. high-elevation species), were filtered at a finer spatial resolution of 1 km^2^ and more widely distributed *Asclepias* species were filtered at a 30-km^2^ resolution. An additional two *Asclepias* species, *A. viridiflora* and *A. curassavica*, were discarded because their records were clearly subject to sample biases and spatial thinning decreased their number of records to less than 40. The process of removing incorrect records and rare species, and spatial filtering, resulted in 24 *Asclepias* species databases each with a minimum of 40 records, totaling 3,549 *Asclepias* records (Table 2).

**Table 2.** Milkweeds models summary and estimated habitat suitability

| *Asclepias* species | Spatial filtering (Km<sup>2</sup>) | Records |  | AUC best model | Estimated Habitat (Km<sup>2</sup>) |  | Host quality weight<sup>1</sup> |
| --- | --- | --- | --- | --- | --- | --- | --- |
|  |  | Total | Filtered |  | Current | Projected |  |
| *A. albicans* | 1 | 434 | 209 | 0.977 | 37,937 | 28,444 | **0.554** |
| *A. asperula* * | 30 | 586 | 244 | 0.815 | 614,502 | 678,420 | 0.078 |
| *A. californica* * | 1 | 607 | 277 | 0.945 | 84,794 | 124,187 | 0.306 |
| *A. cordifolia* * | 30 | 683 | 173 | 0.848 | 230,690 | 215,368 | 0.621 |
| *A. cryptoceras* | 30 | 210 | 128 | 0.839 | 416,872 | 246,545 | 0.218 |
| *A. engelmanniana* * | 30 | 90 | 70 | 0.915 | 277,368 | 230,811 | 0.502 |
| *A. eriocarpa* | 30 | 878 | 145 | 0.888 | 147,561 | 235,229 | 0.541 |
| *A. erosa* * | 30 | 524 | 145 | 0.885 | 285,506 | 466,099 | 0.671 |
| *A. fascicularis* * | 30 | 2259 | 399 | 0.859 | 306,140 | 512,078 | 0.545 |
| *A. halli* | 1 | 48 | 48 | 0.915 | 197,755 | 2,896 | 0.898 |
| *A. incarnata* * | 1 | 144 | 94 | 0.805 | 113,426 | 64,869 | 0.823 |
| *A. labriformis* | 1 | 68 | 57 | 0.945 | 22,126 | 4,863 | **0.554** |
| *A. latifolia* * | 30 | 162 | 102 | 0.858 | 415,185 | 412,476 | 0.394 |
| *A. linaria* * | 1 | 127 | 75 | 0.962 | 376,557 | 382,289 | 0.427 |
| *A. macrosperma* | 1 | 71 | 62 | 0.965 | 624,888 | 124,259 | **0.554** |
| *A. macrotis* * | 1 | 46 | 44 | 0.925 | 196,251 | 151,159 | **0.554** |
| *A. nycatginifolia* * | 1 | 154 | 100 | 0.958 | 112,228 | 233,621 | **0.554** |
| *A. pumila* * | 1 | 89 | 73 | 0.985 | 69,502 | 5,581 | 0.660 |
| *A. soloanoana* * | 1 | 132 | 68 | 0.995 | 17,704 | 65,588 | 1.00 |
| *A. speciosa* | 30 | 1359 | 478 | 0.746 | 958,479 | 937,320 | 0.768 |
| *A. subulata* * | 30 | 627 | 114 | 0.927 | 114,507 | 379,000 | 0.606 |
| *A. subverticillata* * | 30 | 545 | 286 | 0.809 | 594,885 | 674,988 | 0.505 |
| *A. tuberosa* * | 30 | 184 | 88 | 0.831 | 305,144 | 470,641 | 0.681 |
| *A. vestita* * | 1 | 159 | 70 | 0.983 | 34,449 | 80,731 | 0.286 |

\*Species marked with an asterisk are projected to shift their distribution to higher latitudes (S1 File.)
<sup>1</sup>Hostplant quality weight determined by the average weight of monarch larvae reared on 24 *Asclepias* species within a 5-day period. Bold numbers indicate the average weight assigned to species with missing hostplant quality information.

Spatially filtered data were randomly split into training and test data by withholding 25% of the occurrences and the remaining 75% was used for model training. To select background points, we first determined the *Asclepias* environmental domain, corresponding to an area of 50 km^2^ surrounding *Asclepias* occurrences. The environmental domain was then divided into 1 km^2^ grid cells, and background points were randomly selected from within the monarch environmental domain in a checkerboard fashion. Individual *Asclepias* species were modeled using background points from the environmental domain represented by all *Asclepias* species. This process yielded 9000 background points to model *Asclepias* species. The best-fitted models for *Asclepias* with the highest AUC score were used to estimate their current and projected distribution under climate change.

#### Monarch models

The monarch distribution was modeled using a similar approach to *Asclepias*. As described above, we used spatial filtering to correct for sampling biases. Monarch larval records were first filtered at a range of resolutions (1 to 55 km^2^) and the spatial resolution yielding the highest AUC was then selected. The final dataset used to model monarch breeding range was thinned using 30 km^2^ grid cells (the best-fitted model) and included 110 observations. As with *Asclepias*, we withhold 25% of the data for model testing and the remaining 75% was used for model training. To determined monarch larvae environmental domain, we selected 4,000 background points following the same procedure described in the *Asclepias* modeling section, although the number of background points was lower due to the more restricted distribution of monarchs.

To test for the importance of hostplant availability, we first summarized the resulting individual *Asclepias* distribution layers into a single predictor layer representing overall *Asclepias* distribution under current and projected environmental conditions (Fig 1, A and B). The values assigned to grid cell in the genus-level hostplant distribution layer were determined by:

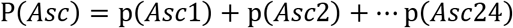

**Figure 1.**
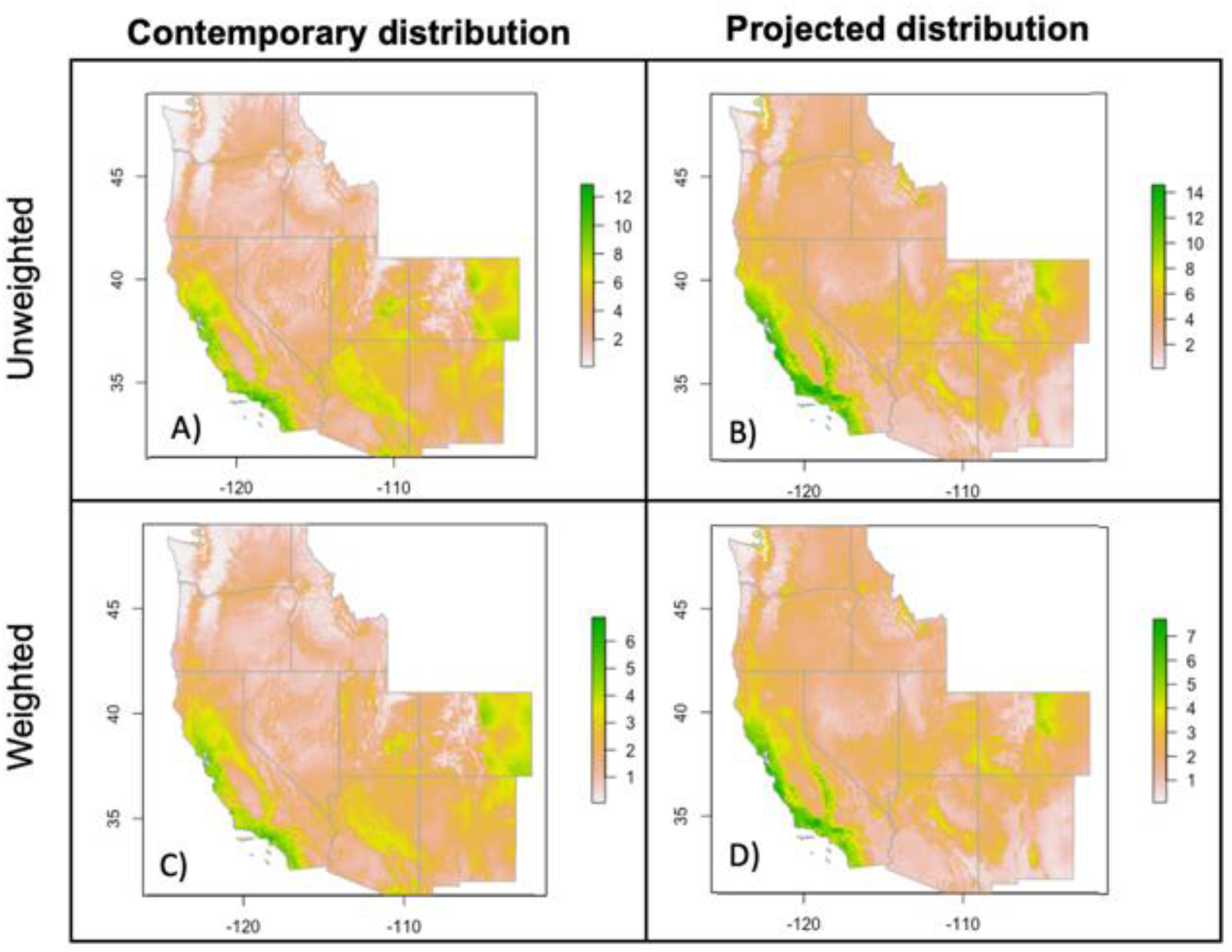
Contemporary and projected genus-level milkweeds distribution. Genus-level *Asclepias* predictor and projection layers used in the monarch hostplant distribution and quality models. The unweighted layers reflect the summed probability of occurrence of all *Asclepias* species, thus reflecting mean probability of occurrence and species richness, while the weighted layer additionally weighs each species by hostplant quality. ‘Green’ indicate high probability of distribution, species richness and/or host-quality, and ‘white’ low probability of distribution, species richness and/or low quality.

Where ‘P’ represents the summarized genus-level probability and ‘p’ probability of distribution of individual *Asclepias* species numbered from 1 to 24. Because the ranges of many *Asclepias* species overlapped, forming the *Asclepias* distribution layer by summing probabilities captures not only the mean probability of distribution but also reflects species richness. We choose this approach under the assumption that higher species richness is associated with increased milkweed abundance and thus higher habitat quality for monarchs. Although a direct assessment of milkweed abundance would be preferable, no such data is readily available. This process is mathematically equivalent to averaging species probabilities and then multiplying by species richness.

To assess hostplant quality, we used the average monarch larval weight supported by each *Asclepias* species grown under greenhouse conditions (Table 2). These protocols are described in detail by Petschenka and Agrawal (Petschenka and Agrawal 2015). Briefly, *Asclepias* plants were grown from seed in a greenhouse and after a period of 4-7 weeks neonate monarch caterpillars were placed individually upon the leaves of potted plants and weighed after 5 days. Assessing hostplant quality under controlled greenhouse conditions controls for extraneous factors such as natural predator, competition with other herbivores, induced plant defenses and environmental variation that are necessarily associated with a field bioassay.

We weighted each *Asclepias* species distribution layer according to its host quality. The Milkweed with the greatest larval weight (*A. sololana*) was given a value of 1, and all other species were assigned values as proportions of this value, with the lowest quality weight being 0.078 (*A. asperula*) (Table 2). Five species with no information on larval weight were weighted by the average host quality weight of 0.55. Weighted layers were then summarized into a single predictive layer representing the hostplant probability of distribution and species richness weighted by hostplant quality (Fig 1, C and D). The values assigned to grid cells of the overall hostplant quality layer were calculated as follow:

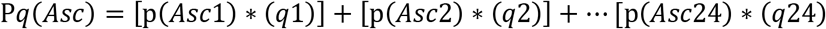

Where ‘P*q*’ represents the summarized genus-level probability of distribution weighted by hostplant quality, ‘p’ the probability of distribution of individual *Asclepias* species numbered from 1 to 24, and ‘*q*’ the host-quality weight estimated for each *Asclepias* species. This approach is parallel to that used for the *Asclepias* distribution layer (above), capturing the effects of both the mean probability of distribution and species richness for all co-existing *Asclepias* species, but now weighting each species according to its relative hostplant quality.

Lastly, we used the summarized *Asclepias* layers as predictors to generate and compared three models: a null model (climate envelope model) using only environmental factors as predictors; a model using environmental factors and hostplant distribution represented by the summarized *Asclepias* distribution (hostplant-presence model); and a second model using environmental factors and *Asclepias* distribution weighted by host quality (hostplant-quality model).

To identify the variables contributing more to each model, in addition to the “Analysis of Variable Contribution” reported by MaxEnt, we performed a jackknife test of variable importance. In a jackknife test, models are re-run using a single variable in isolation to identify the variables that yield the highest model gain when used in isolation. This test also identifies those variables that, when removed, decrease the model gain the most by re-running the models excluding one variable at a time.

Since there is currently no consensus regarding a single most appropriate metric to evaluate SDMs performance (Peterson et al. 2008, 2011, Warren and Seifert 2011), we evaluated model performance based on several criteria. The area-under-the-curve (AUC) statistic provides an estimate for the accuracy of predictions, with 0 indicating no predictive accuracy and 1 perfect predictive accuracy. An AUC score of 0.5 indicates that the model performs no better than random. We also estimated performance metrics based on the Akaike Information Criterion corrected for small sample sizes (AICc). The AICc metric have the advantage of balancing both, model goodness-of-fit and model complexity. Furthermore, compared to AUC and BIC (Bayesian-Information-Criterion) based methods, AICc evaluation methods have been shown to favor models that more accurately estimate the relative importance of variables and habitat suitability, both in the training region and when models are extrapolated to a different time period (Warren and Seifert 2011). We calculated the AICc, delta AICc (ΔAICc), and Akaike weights (wAICc) for each model using the ENMeval package (Muscarella et al. 2014). The model with the lowest AICc value is considered the best model out of various candidate of models. The ΔAICc is the difference between the best AICc and other candidate models. The best candidate model has a ΔAICc of 0 and models with ΔAICc lower than 2 are generally considered to have substantial support and should not be discarded (Muscarella et al. 2014). Akaike weights (wAICc) represents the likelihood of a model given the data. The weights are normalized to sum 1 and are interpreted as probabilities (Burnham and Anderson 2004).

Finally, we estimated suitable breeding area for monarchs and for *Asclepias* distribution from polygons drawn around areas with grid cell values higher than 0.5 from the output logistic layers projected from the final models.

## Results

### Asclepias models and estimated distribution

All *Asclepias* final models had AUC scores higher than 0.8, except for *A. speciosa* model which yielded an AUC score of 0.74, indicating that these models are a good fit for the observations (Table 2). The current estimated distributional ranges (Appendix S1, left panels.) were consistent with *Asclepias* spp. distributions published by the Biota of North America Program (BONAP) (Kartesz 2015).

Overall, within the study area, half of the *Asclepias* species are projected to expand their ranges by a mean of 88% (i.e. nearly doubling their distributions) whereas the other half will contract their ranges by a mean of 42% (i.e. more than halving their distributions) (Table 2 and Appendix S1). Of the 24 *Asclepias* species, 19 species are predicted to shift their distributions to higher latitudes (79%) both along the Pacific coast and inland, with 11 of these also expanding their distributions. Of the 4 species not shifting their distributions northward, 3 will contract their ranges.

### Monarch models and estimated distribution

The AUC scores did not differ considerably among the three models, but AUC values were slightly higher for the hostplant-presence model (0.803) compared to both the hostplant-quality (0.800) and climate envelope model (0.799). However, the AIC-based metrics preferred the hostplant-presence model (ΔAICc=0, wAICc=1.00) over the hostplant-quality (ΔAICc=123.50, wAICc=1.515^-27^) and climate envelope models (ΔAICc=168.52, wAICc=2.545^-37^). The ΔAICc for the competing climate envelope and hostplant quality model was much larger than 2 indicating that these two models had limited support. Likewise, the wAICc of the hostplant model was nearly 1 suggesting that the likelihood of this model being the best-fitted model was high (Table 3).

**Table 3.** Monarch model performance comparison and estimated habitat suitability

| Model | $n^1$ | AUC | AICc | $\Delta AICc$ | wAICc | Estimated habitat (km<sup>2</sup>) * | |
| --- | --- | --- | --- | --- | --- | --- | --- |
|  |  |  |  |  |  | Current | Projected (2070) |
| Climate envelope | 60 | 0.799 | 25,694.99 | 168.52 | 2.545<sup>-37</sup> | 214,245 | 409,091 |
| Hostplant-presence | 61 | 0.803 | 25,526.46 | 0.000 | 1.00 | 252,464 | 466,306 |
| Hostplant-quality | 61 | 0.800 | 25,649.97 | 123.50 | 1.515<sup>-27</sup> | 252,465 | 486,200 |

\*Estimated habitat was calculated by summarizing areas with probability of distribution higher than 0.5 from the logistic output layers produced by each model. <sup>1</sup> $n$ gives the number of parameters of each model.

The environmental variables that contributed the most to the climate envelope model were the “minimum temperature of the coldest month” (43.4% contribution; (Fick and Hijmans 2017) and “precipitation seasonality” (25.2% contribution, Fig. 3 A; (Fick and Hijmans 2017)). For both hostplant-presence and hostplant quality models, the hostplants variable was the second most important factor for predicting the western monarch breeding range. The hostplants variable contributed most to the hostplant distribution model (22.5%), after the “minimum temperature of the coldest month” (33.1%) (Fig. 3 B and C). Although the hostplant quality model did not produce the best-fit model, weighting the hostplant layer by host-quality increased the contribution of the hostplant variable by 3% and decreased “minimum temperature of the coldest month” variable contribution by 7% compared to the hostplant distribution model (Fig 3, B and C). Both hostplants layers (weighted by host-quality and unweighted) exhibited the highest gain (>0.40) in the jackknife test for variable importance in both hostplant models (Appendix S2). This indicates that hostplants provided the most useful information for predicting where monarch breeding grounds occur. For all three models the “average precipitation of the warmest quarter” (Fick and Hijmans 2017) decreased model gain the most when omitted suggesting that this environmental variable has the most information that is not present in other variables (Appendix S2).

The process of weighting the *Asclepias* distribution layer by quality did not dramatically altered the hostplant layer, and mainly rescaled the values of the layer (Fig 1, lower panels). This was probably due to large range overlaps among *Asclepias* species as it can be observed by overlaying the polygons corresponding to *Asclepias* suitable habitat (Appendix S3). Thus, a grid cell occupied by multiple *Asclepias* with variable host-quality may have the same value as a grid cell occupied by a few high-quality *Asclepias* species. The only area where weighting hostplants by quality appeared to change the grid cell values of the hostplant quality layer was the southwest region of Arizona and Utah which appeared to be occupied mostly by lower quality species, predominantly by *A. asperula*, our lowest quality hostplant (Fig 1, lower panels and Table 2).

The climate envelope model estimated more restricted ranges for the contemporary and future monarch distributions. Both hostplants models estimated nearly identical contemporary distributions for monarchs that were ~18% larger than the estimated by the climate envelope model (Fig 2, left panels and Table 2). Although all three models predicted future range expansions that nearly doubled their corresponding contemporary estimates, the hostplant-presence and hostplant-quality models projected an increased in habitat suitability 14 and 19% larger than that of the climate envelope model, respectively (Fig 2, right panels and Table 2). This difference was primarily due to the fact that the climate envelope model predicted range contractions inland whereas both hostplants models predicted range expansions in this region. Finally, we detected some slight differences in the areas where hostplants models predicted that such range expansions will occur. For example, the hostplant quality model predicted a smaller range for monarchs in western New Mexico and a larger range in central Nevada, Utah and western Colorado. (Fig 2, right panels).

**Figure 2.**
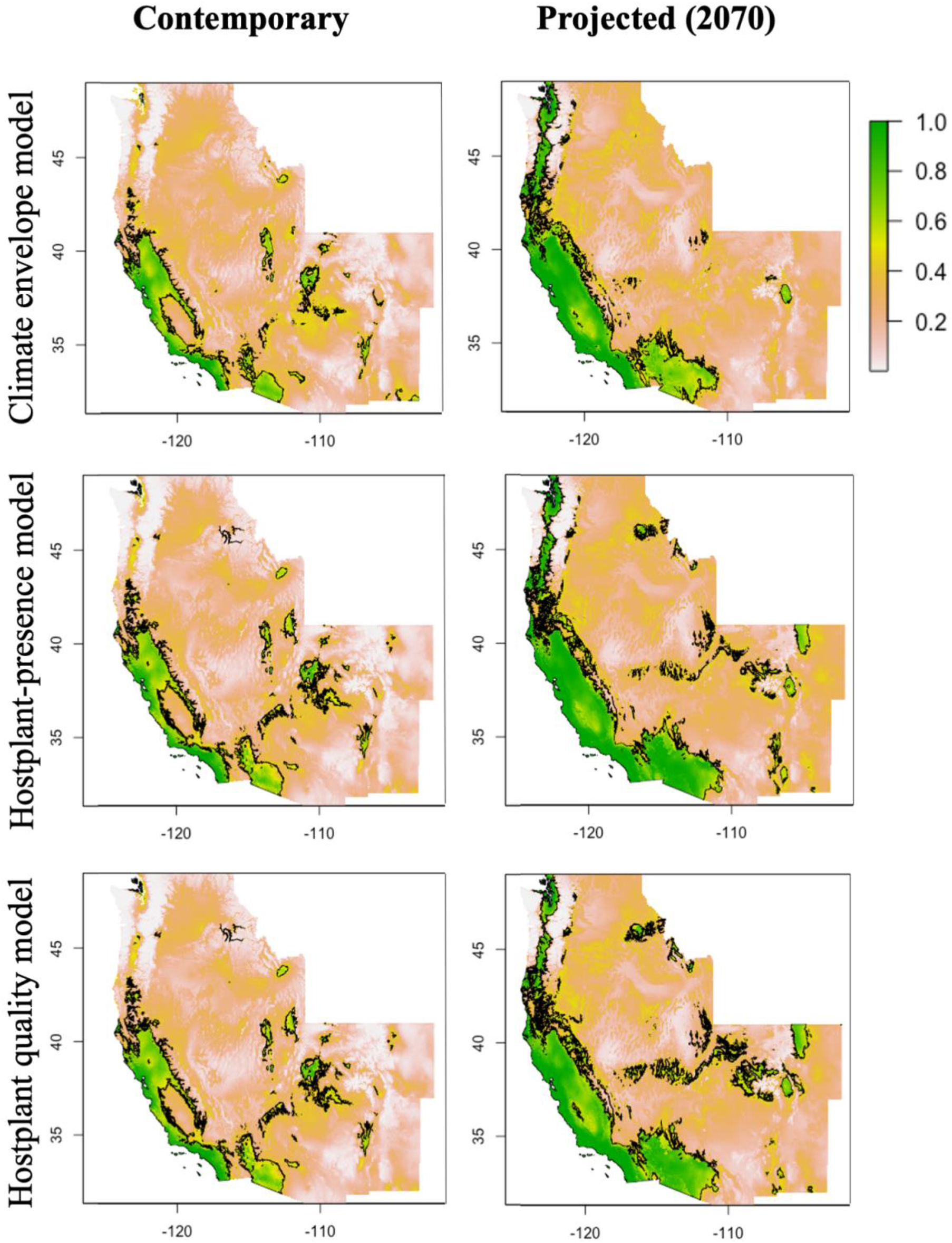
Monarch contemporary and projected future breeding ranges. Left panels (A, C, E) indicate the current probability of distribution of the monarch breeding grounds estimated by the three models, with ‘light yellow’ representing low probability and ‘dark blue’ high probability. Right panels (B, D, F) indicate the projected probability of distribution of the monarch breeding grounds for the year 2070 estimated by the three models. Suitable habitat for monarch breeding is delineated in black and represent areas with a probability of distribution greater than 0.5.

**Figure 3.**
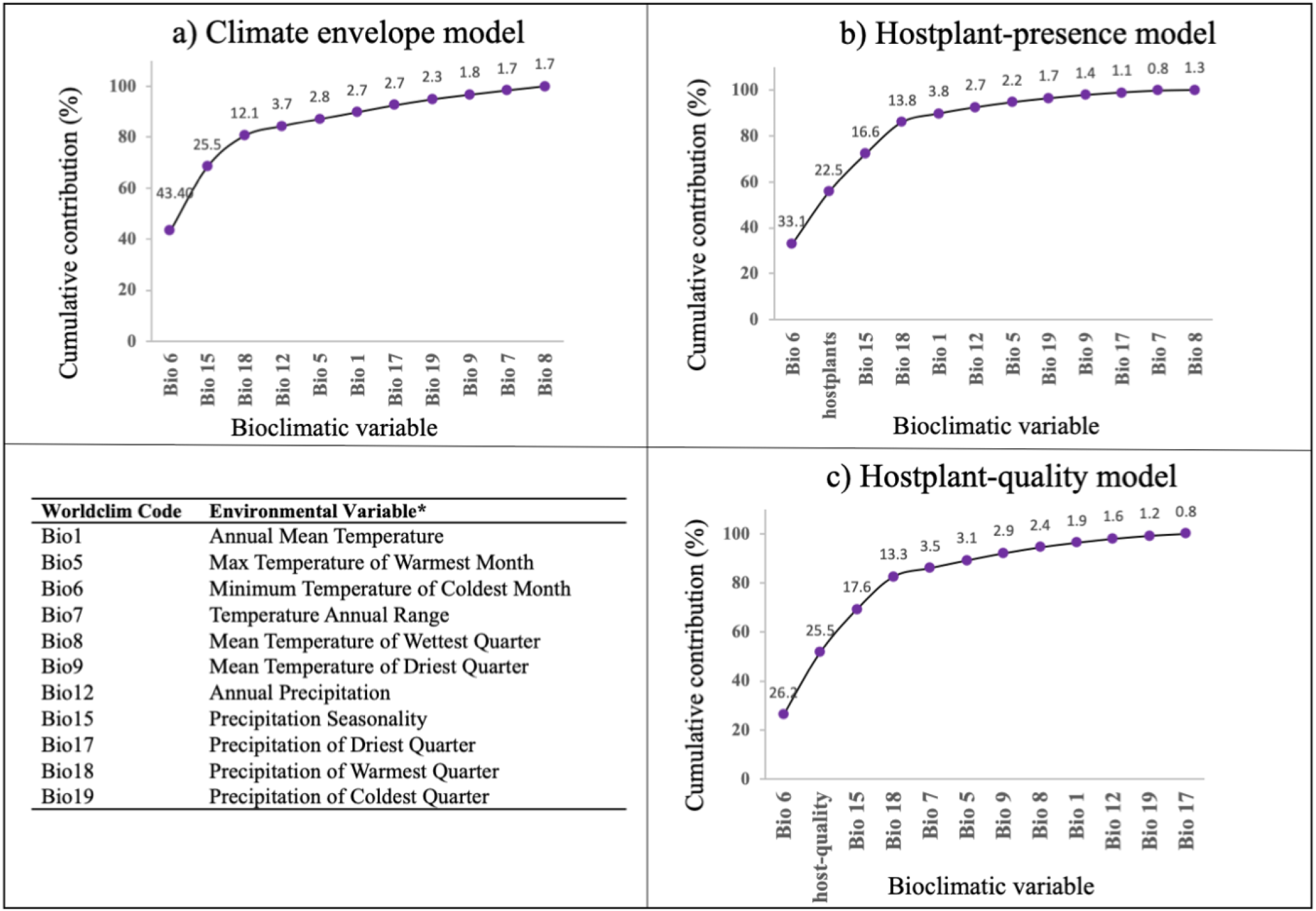
Analysis of variable importance. Percent contribution that each variable contributes to the models in decreasing order from left to right.

## Discussion

Predicting herbivore response to climate change requires incorporating future hostplant availability, but hostplant quality may play a secondary role. While climate envelope projected a more restrictive current monarch distributions than hostplant models, model comparisons suggested that hostplant information provided superior predictive power. Furthermore, the three models differed in their future monarch projections under climate change with models including hostplant information predicting an increased in habitat suitability 14-19% larger than that of the climate envelope model. Despite the importance of hostplant information, models including hostplant quality did not prove superior to the model based on hostplant presence. Our study suggests that information on critical biotic interactions is essential to predict future species distributions under climate change.

The hostplant model was preferred by AIC-based metrics over a traditional climate envelope model and hostplant quality model. Hostplant availability, together with the minimum temperature of the coldest month, contributed over fifty percent to the hostplant model and over forty percent to the model gain when used in isolation. This suggests that the western monarch breeding ranges are co-limited by both cold temperatures and hostplant availability. These findings are consistent with past work by Lemoine (Lemoine 2015) who found that models incorporating hostplants and environmental factors most accurately estimated the eastern monarch distribution. Additionally, our results are also supported by previous studies from Steven and Frey (Stevens and Frey 2010) and Dilts *et al*. (Dilts et al. n.d.) who identified *Asclepias* availability as well as climatic variables, including minimum temperature of the coldest month, as key for structuring the western monarch breeding grounds. These findings add to the increasing body of evidence suggesting that biotic interactions may govern species distributions as strongly as environmental conditions (Dilts et al. n.d., Araújo and Luoto 2007, Preston et al. 2008, Schweiger et al. 2008, de Araújo et al. 2014, Fraterrigo et al. 2014, Lemoine 2015, da Cunha et al. 2018).

Hostplant quality varied ten-fold among Milkweed species but did not have a large effect on the overall estimates for the contemporary distribution of monarchs. We speculate this result may be due to the fact that our genus-level distribution layer also reflected species richness. *Asclepias* ranges exhibit substantial range overlaps in the American West (Appendix S3), thus, adult monarchs may preferentially oviposit on higher quality milkweeds in areas with mixed quality resources (Gripenberg et al. 2010), diminishing the influence of low-quality species. Hostplants model projected similar monarch distributions under a climate change scenario; however, their projections differed in some regions of the inland states of Utah, Nevada, New Mexico and Colorado. This implies that the importance of hostplant quality in determining herbivore distributions should not be discarded altogether as it may play a significant role in instances where herbivores rely on hostplant with less geographic overlap, and therefore, fewer food choices.

Our results demonstrate how climate envelope models that accurately represent current distributions may provide poor prediction for the future. This can occur when critical distributional drivers (e.g. hostplant distributions) correlate strongly with environmental factors under contemporary conditions (Wharton and Kriticos 2004) but not under climate change. These mechanistically-flawed models thus provide inaccurate predictions (Brewer and Gaston 2003, Soberon and Nakamura 2009). In our study, the climate envelope model– although more restricted estimated very similar monarch contemporary ranges than hostplant models (Fig 2, left panels) but differed in their future projections (Fig 2, right panels). Specifically, the two hostplant models predicted larger range expansions of monarchs than the climate envelope model inland. This suggests that the climate envelope model over predicted monarch climatic limitations due to contemporary correlations between climatic factors and milkweed distributions, but that this correlation may not persist in the future. Accordingly, models based solely on climatic factors may be adequate for estimating contemporary species distributions but nevertheless produce misleading projections under novel circumstances where abiotic conditions and biotic interactions do not respond in tandem to climate change.

The importance of incorporating the climatic response of hostplants into models is underscore by the fact that only models including hostplant information predicted range expansion inland while the climate-envelope model did not (Fig 2, right panels). The predicted inland range expansions of the western monarch breeding range appeared to be driven by higher hostplant availability in the regions of central Nevada, Utah and Colorado under future climatic conditions, which was identified by our models as one of the most important factors delimiting monarch distributions. Our results are congruent with previous findings by Lemoine (Lemoine 2015) whose study predicted northern range expansion of the eastern monarch population resulting from projected *Asclepias* range expansions under future climate change scenarios.

Lastly, it is worth noting that our model projections do not consider factors that were beyond the scope of our study but that may significantly impact monarch future distributions. For example, pesticide and land-use practices, specially overwintering habitat loss to housing development, is an existing threat to monarchs habitat (Pelton et al. 2019) that is likely to persist in the upcoming years. Furthermore, dams and human-facilitated invasions, may alter riparian areas potentially disrupting monarch migration patterns and monarch breeding grounds. Autumn migrants often follow riparian corridors (Dingle et al. 2005) and riparian vegetation has been associated with habitat suitability for some western Milkweed species (*A. subulata*, and *A. asperula*) (Dilts et al. n.d.).

## Conclusions

In summary, this study shows that accounting for biotic interactions– and their distributional response to climate change– is required to predict the future distributions of species obligately dependent on such interactions. A climate-envelope approach may be effective for estimating contemporary species distributions but may produce misleading future projections as climate change may uncouple suitable climate from essential biotic interactions. Hostplant quality did not play a significant role in delimiting monarch distribution in the American West where *Asclepias* ranges overlapped substantially. However, there were slight differences in some regions suggesting that host-quality may still be important for predicting distributions of species dependent on a fewer number of resources. These results are relevant, not only for most herbivorous insects which are highly host-specific, but also for all organisms incurring in obligate biotic interactions (e.g. parasitic or mutualistic interactions). Ultimately, accurate projections for the future will require better incorporating inter-specific dynamics into our models.

## Acknowledgements

We thank Anurag Agrawal for providing hostplant quality data and Stijn Hantson, Colleen Neil, and Will Petry for their advice on the data collection and modeling phase of this project. We also thank the Monarch Larvae Monitoring Project (MLMP) for kindly providing monarch larval records.

## Appendices

**Appendix S1. Milkweeds species contemporary and projected distributions**. Estimated contemporary distribution of 24 species of *Asclepias* (left panels) and projections under a climate change scenario for 2070 (right panels), with ‘green’ indicating high probability of distribution and ‘white’ low probability. Suitable habitat for each *Asclepias* spp. is delineated in black and represent areas with a probability of distribution greater than 0.5.

**Appendix S2. Jackknife Test of Variable Importance.** Jackknife test of variable importance. Blue bars indicate model gain when each variable is used in isolation, turquoise bars represent model gain when a single variable is excluded, and red bars represent model gain when all variables are included.

**Appendix S3. Milkweeds range overlaps in the Western United States.** Each overlaid layer represents the range of individual *Asclepias* species estimated by drawing a polygon around areas with areas with a probability of distribution greater than 0.5. Regions in white represent areas with no Milkweeds, ‘light green’ represent low range overlap, and ‘dark green’ high range overlap.

**Metadata S1.** R code for monarch and milkweed species distributions modeling.

**Data S1. Milkweed species records** retrieved from various open source databases within the study region.

**Data S2. Monarch larval records** retrieved from open source databases and the MLMP (Monarch Larvae Monitoring Project) within the study region.

