## Supplementary figures and images for "Roles of hostplant availability and quality for the distribution and climate change response of a dietary specialist herbivore"

### Supplemental Figure 1

#### Milkweeds Contemporary and Projected Distributions

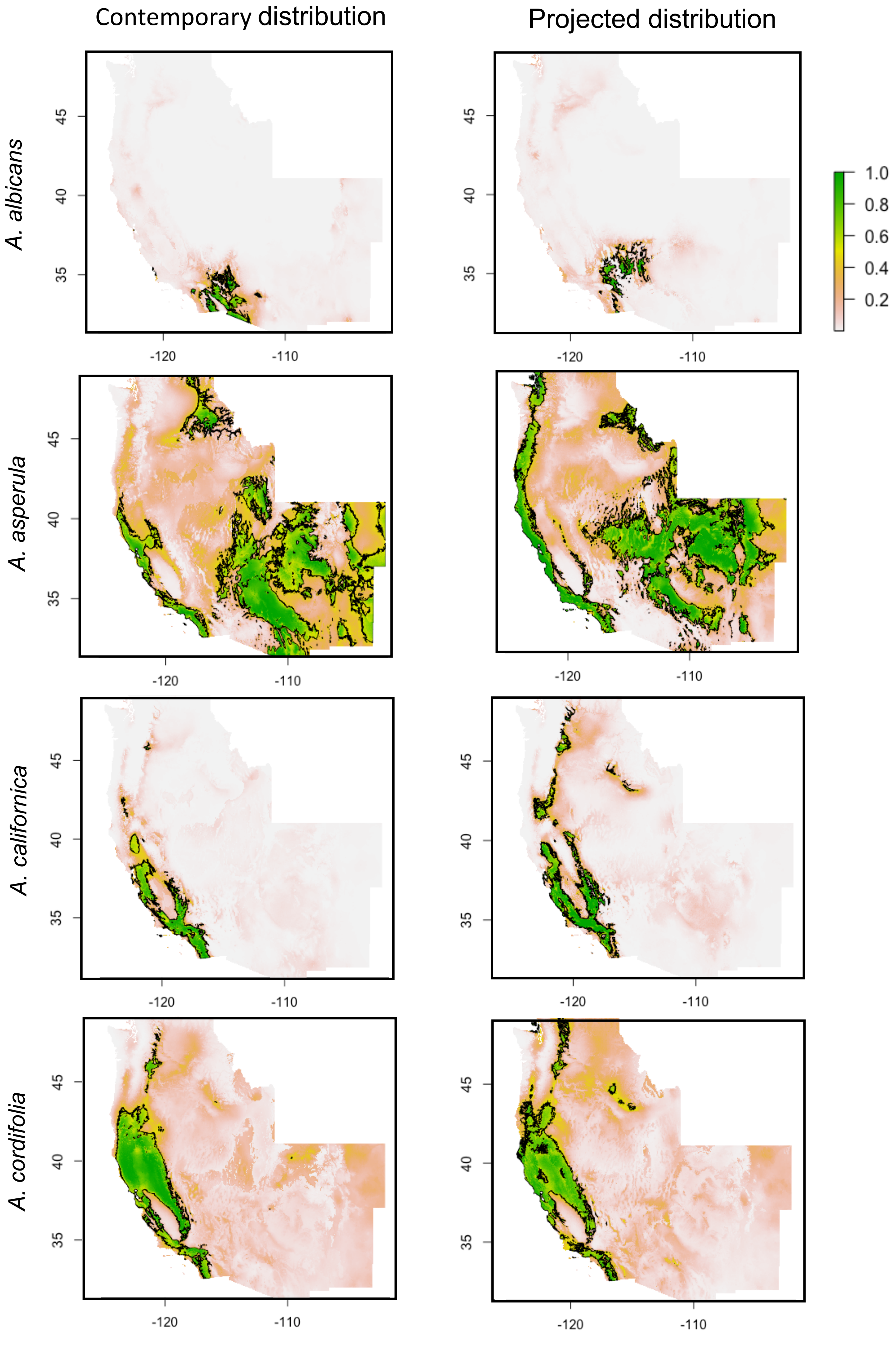

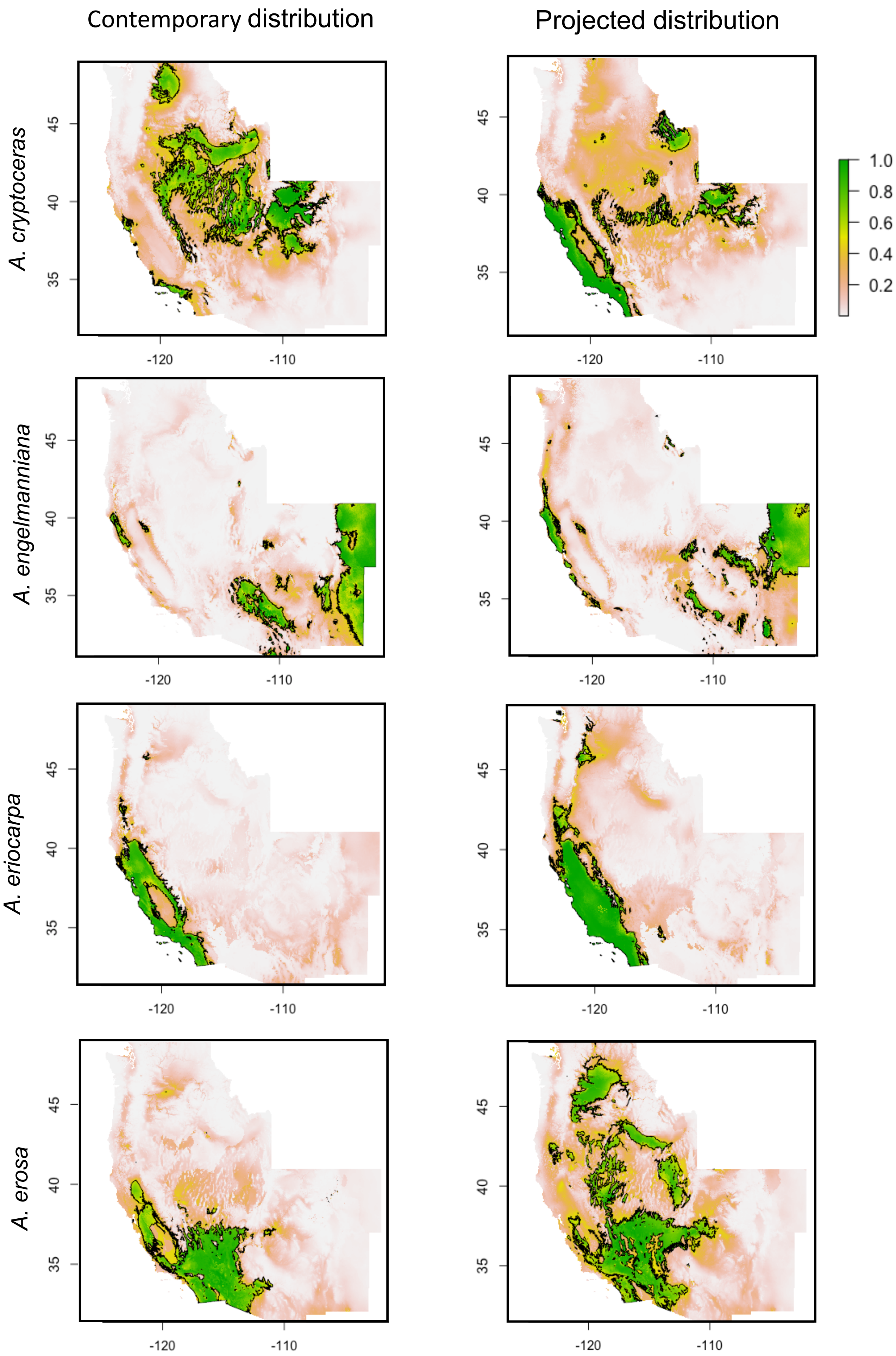

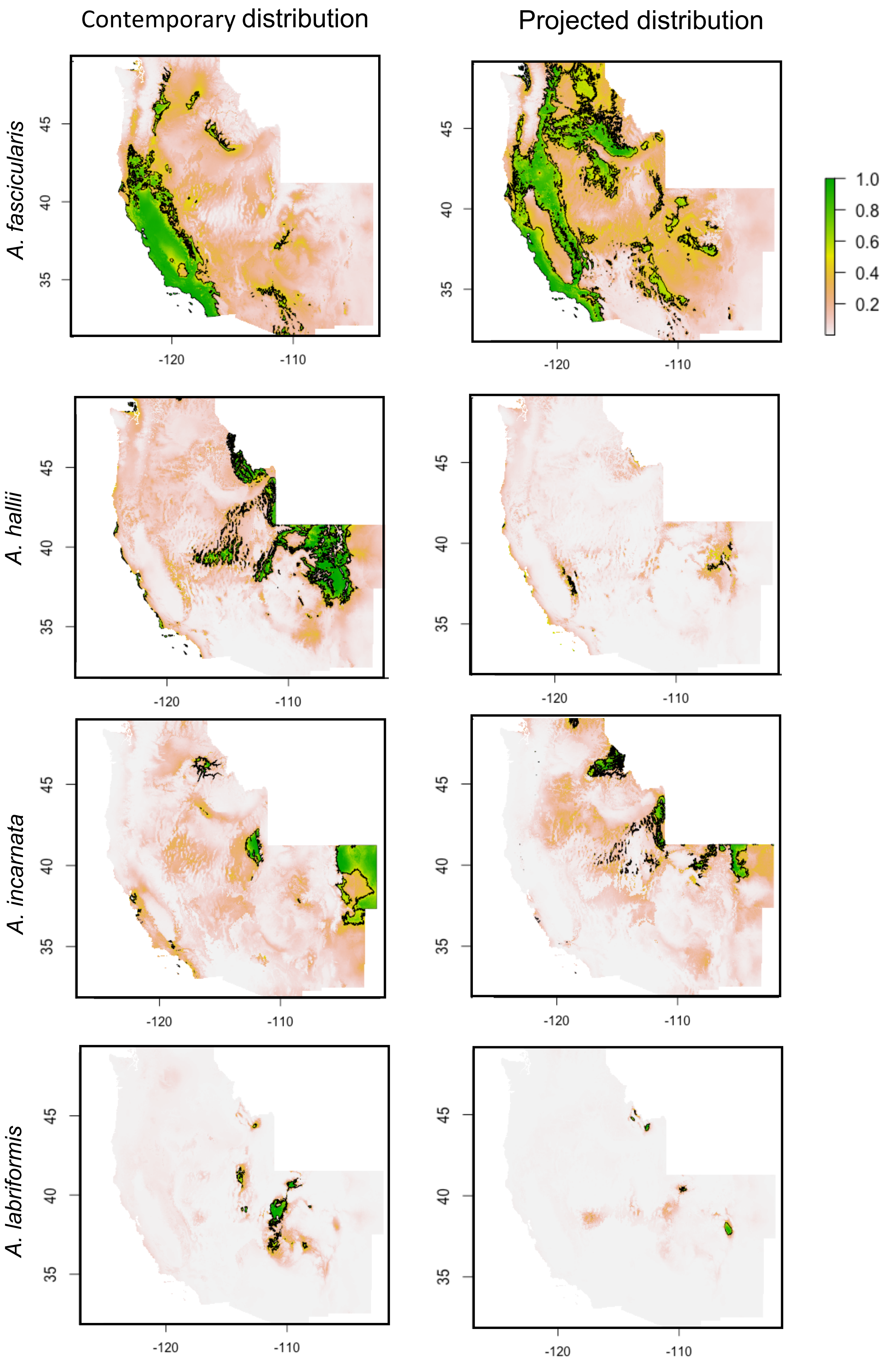

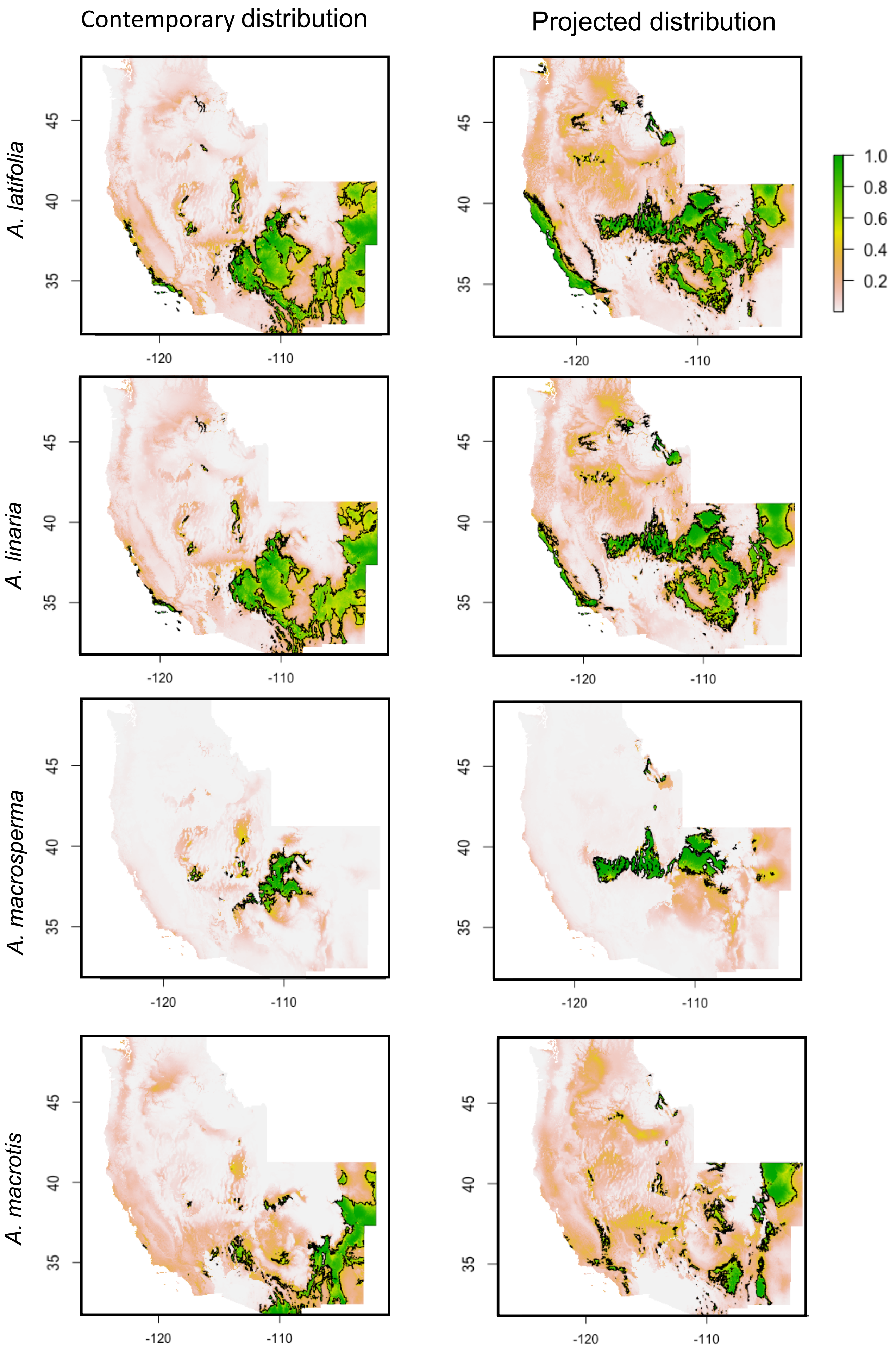

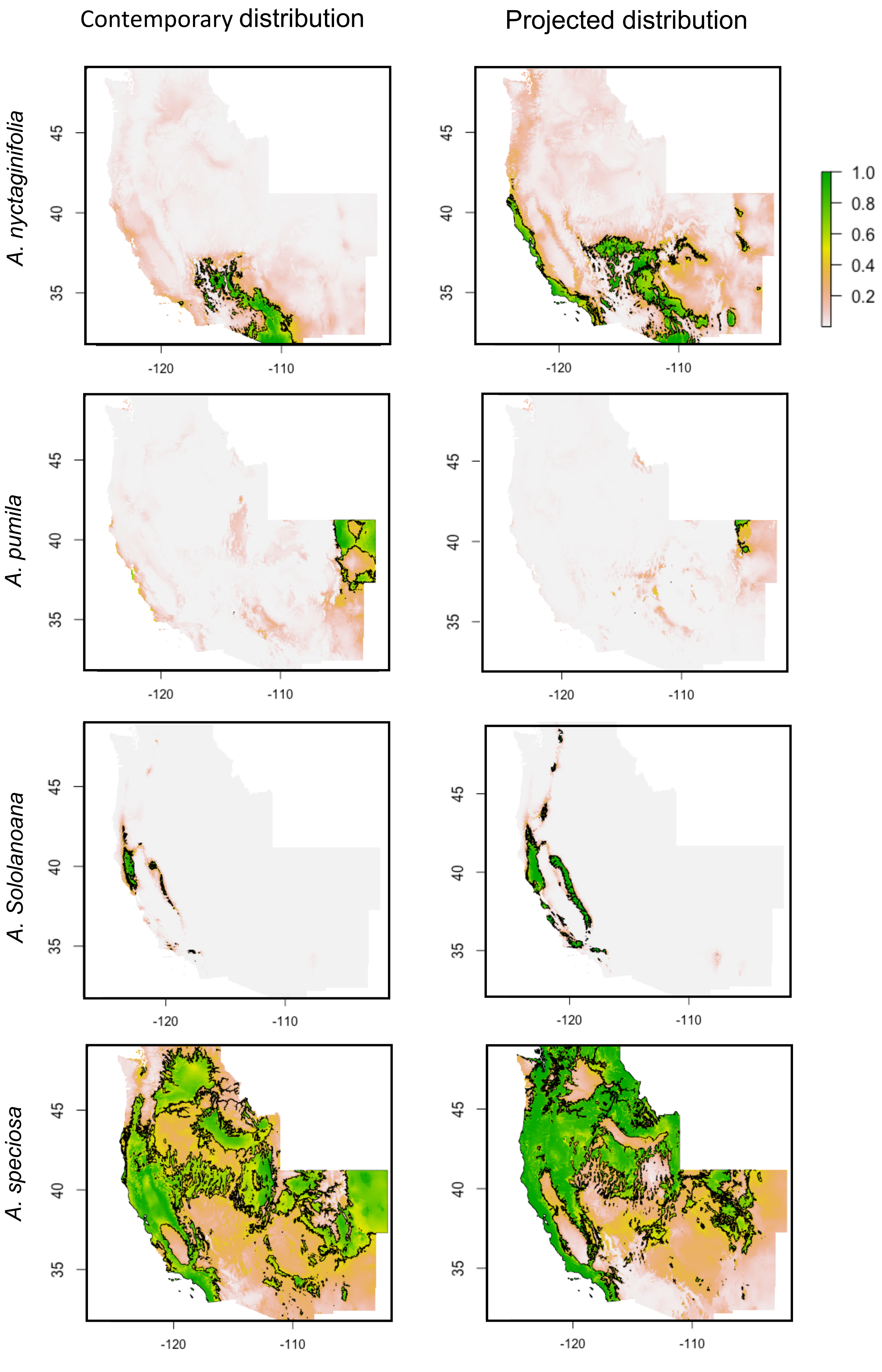

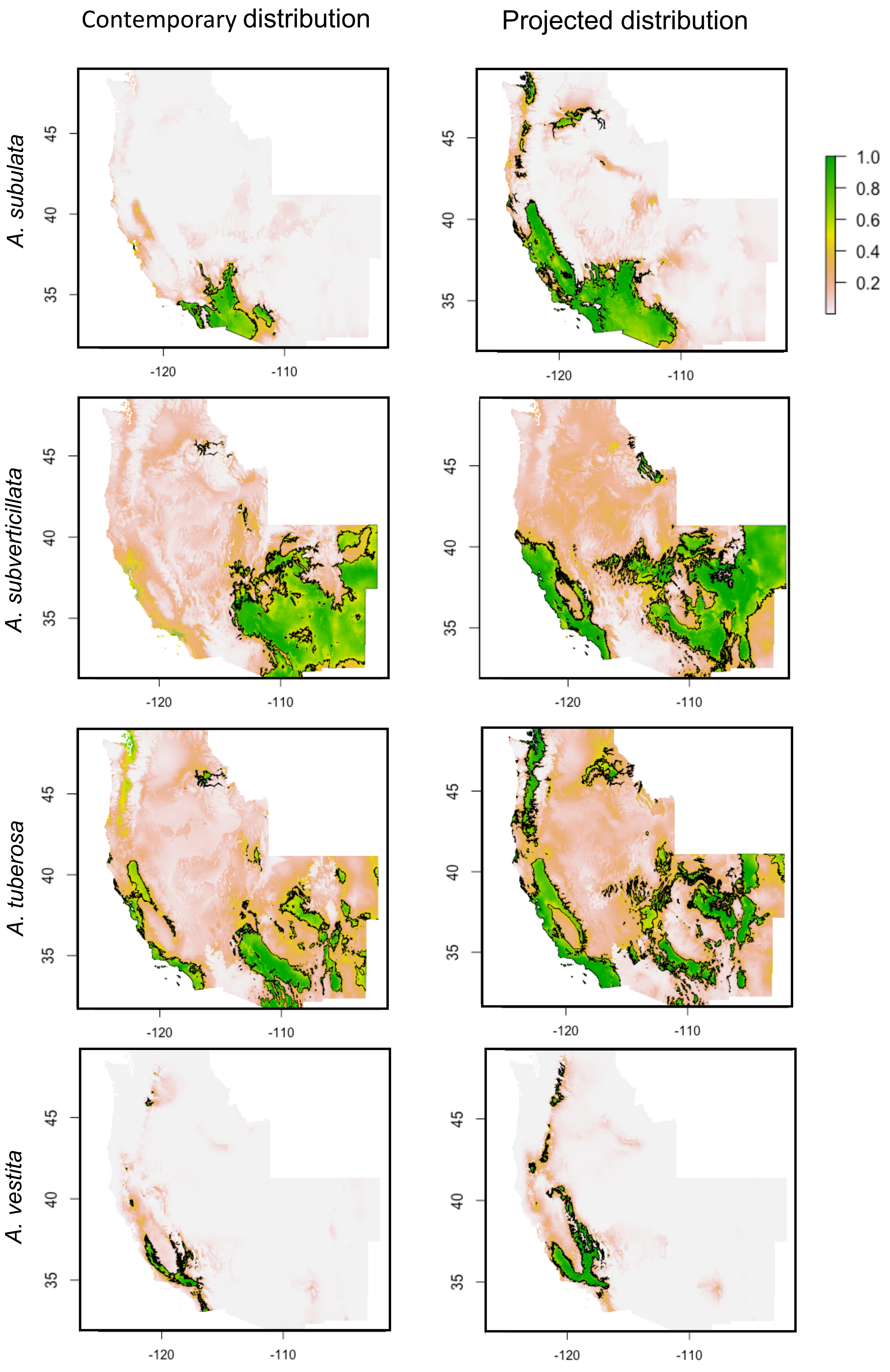

### Supplemental Figure 3

#### Milkweed ranges overlap in the western United States

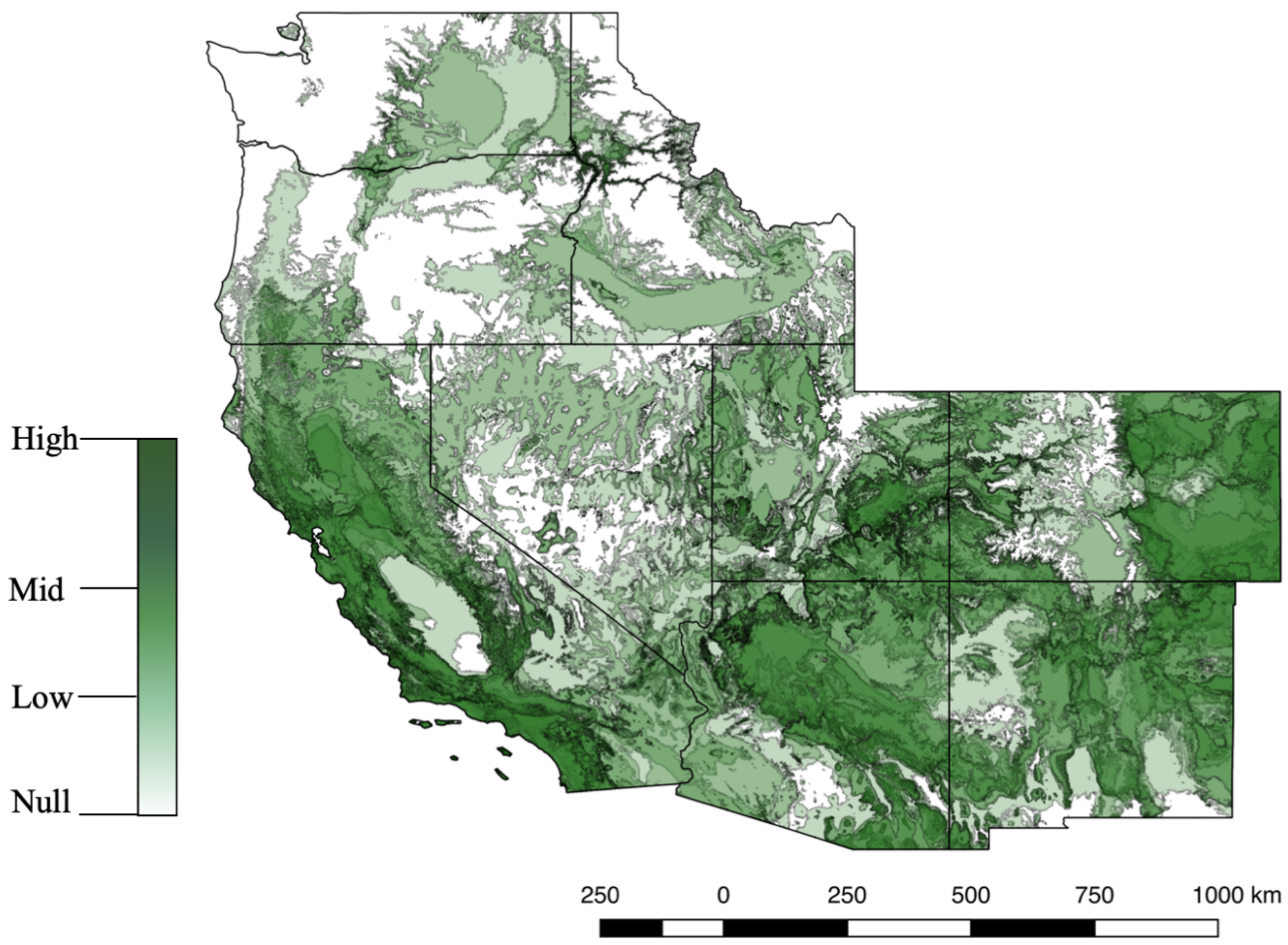
