## Supplemental Figure 2 for "Roles of hostplant availability and quality for the distribution and climate change response of a dietary specialist herbivore"

### Jackknife test of regularized training gain

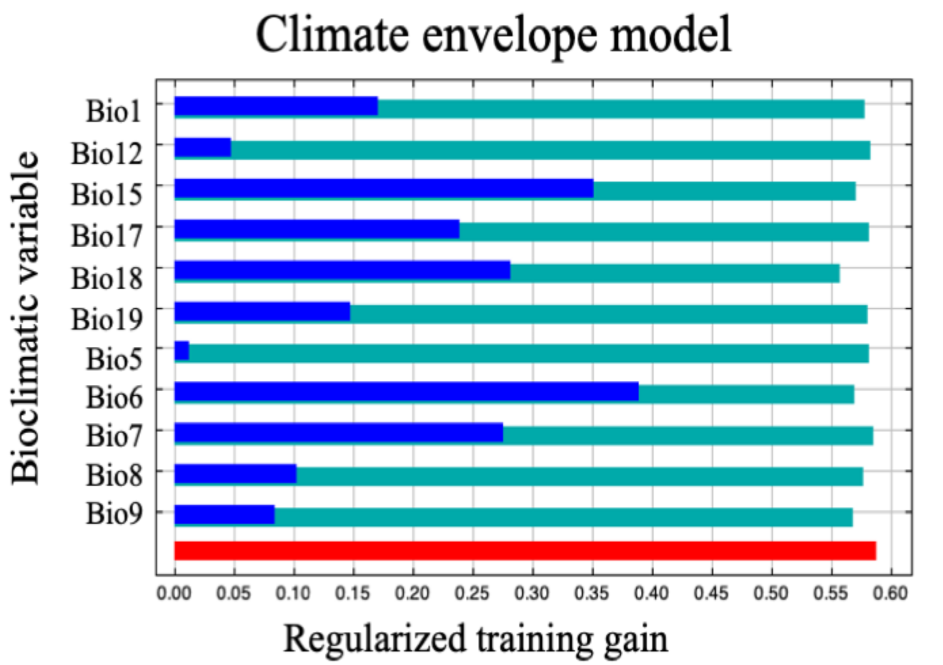


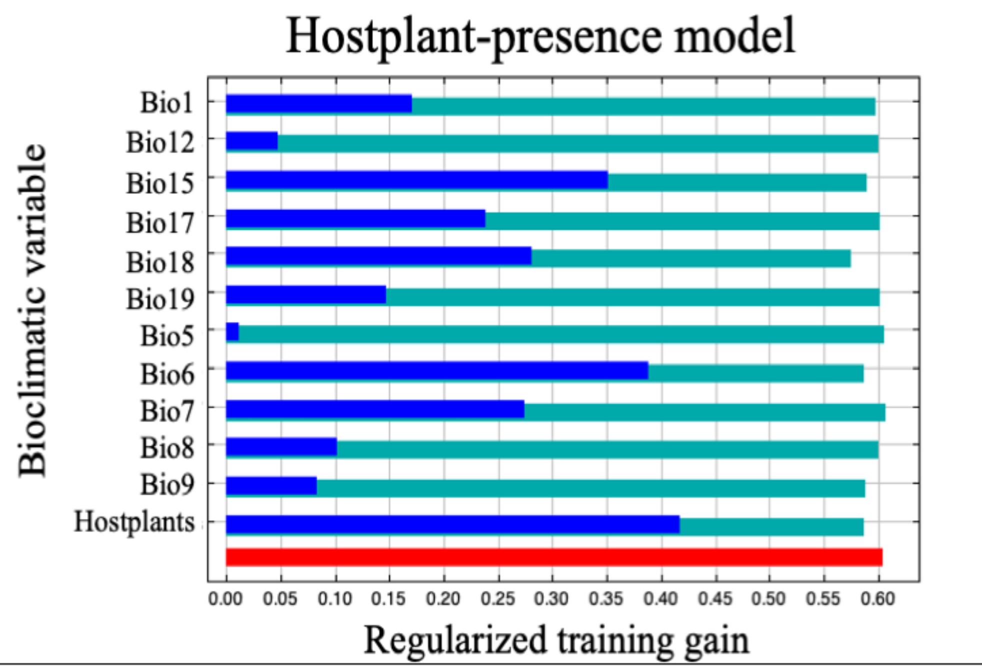


| Worldclim Code | Environmental Variable* |
|---|---|
| Bio1 | Annual Mean Temperature |
| Bio5 | Max Temperature of Warmest Month |
| Bio6 | Minimum Temperature of Coldest Month |
| Bio7 | Temperature Annual Range |
| Bio8 | Mean Temperature of Wettest Quarter |
| Bio9 | Mean Temperature of Driest Quarter |
| Bio12 | Annual Precipitation |
| Bio15 | Precipitation Seasonality |
| Bio17 | Precipitation of Driest Quarter |
| Bio18 | Precipitation of Warmest Quarter |
| Bio19 | Precipitation of Coldest Quarter |

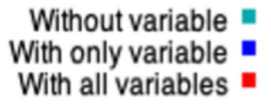


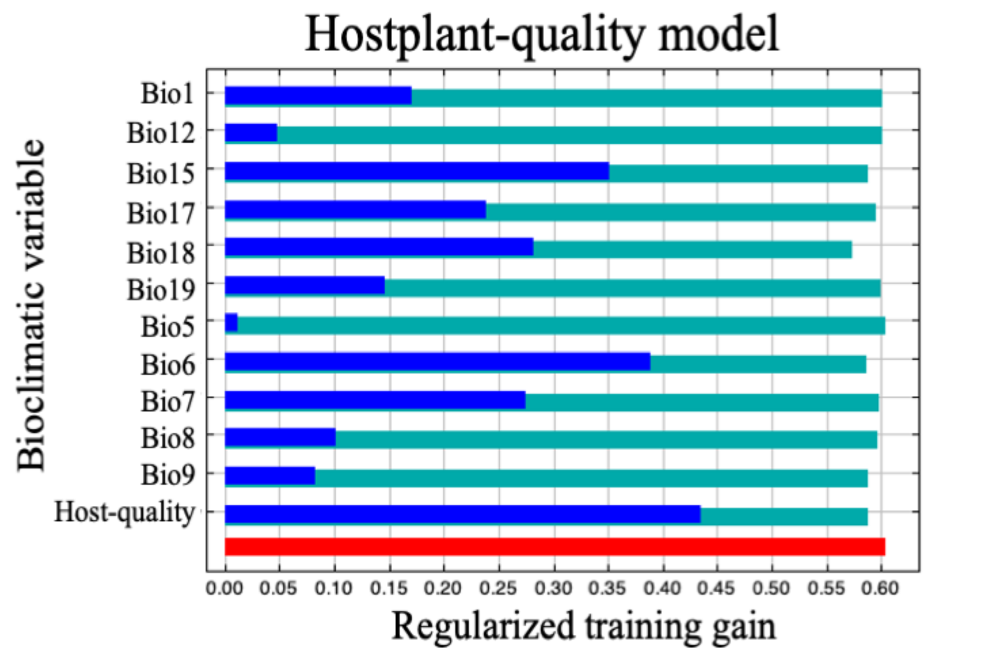
